# Highly specific, multiplexed isothermal pathogen detection with fluorescent aptamer readout

**DOI:** 10.1101/2020.02.18.954719

**Authors:** Lauren M. Aufdembrink, Pavana Khan, Nathaniel J Gaut, Katarzyna P. Adamala, Aaron E. Engelhart

## Abstract

Isothermal, cell-free, synthetic biology-based approaches to pathogen detection leverage the power of tools available in biological systems, such as highly active polymerases compatible with lyophilization, without the complexity inherent to live-cell systems, of which Nucleic Acid Sequence Based Amplification (NASBA) is well known. Despite the reduced complexity associated with cell-free systems, side reactions are a common characteristic of these systems. As a result, these systems often exhibit false positives from reactions lacking an amplicon. Here we show that the inclusion of a DNA duplex lacking a promoter and unassociated with the amplicon, fully suppresses false positives, enabling a suite of fluorescent aptamers to be used as NASBA tags (Apta-NASBA). Apta-NASBA has a 1 pM detection limit and can provide multiplexed, multicolor fluorescent readout. Furthermore, Apta-NASBA can be performed using a variety of equipment, for example a fluorescence microplate reader, a qPCR instrument, or an ultra-low-cost Raspberry Pi-based 3D-printed detection platform employing a cell phone camera module, compatible with field detection.

## Introduction

Low-cost, rapid pathogen detection methods are of high interest for field diagnostics and use in the developing world, both for disease detection, and pathogen monitoring (e.g., in water supplies or agricultural products). Several nucleic acid amplification techniques have been developed to identify disease-causing pathogens, of which the polymerase chain reaction (PCR) is the best-known.^1^ PCR is robust, specific and widely used, and it constitutes the basis for a wide range of diagnostic tests. However, the thermocyclers required to perform PCR are energy intensive and expensive (ca. $5-10K), making them cost-prohibitive for field diagnostics or use in low-resource environments. Furthermore, the typical readout of PCR, electropherograms, requires trained technicians and additional equipment. It is possible to monitor PCR reactions in real time, but this requires the coupling of a fluorescence detection system to the thermocycler, increasing the cost of the instrument by as much as an order of magnitude. These reactions are typically monitored by expensive oligonucleotide probes or non-specific DNA intercalating dyes.^2^

Isothermal nucleic acid amplification techniques are an attractive means to mitigate the costs and complexity associated with thermal cycling. Several isothermal techniques have been reported, such as nucleic acid sequence based amplification (NASBA), loop mediated amplification, recombinase polymerase amplification (RPA), helicase dependent amplification, and SHERLOCK.^3–7^ In each case, these reactions employ strand displacement, nucleases, or recombinases to generate double-stranded DNA, either as a product, or an intermediate that is transcribed to an RNA output. Reactions that generate RNA outputs are of high interest, given that a number of RNA polymerases can readily be overexpressed, lyophilized (negating the need for a cold chain), and used to generate large quantities of RNA in these reactions. Furthermore, unlike dsDNA, RNA can exhibit diverse functional behaviors, including binding of fluorogenic dyes and enzymatic activity, enabling a wide range of potential readouts.^8^

A common pitfall associated with isothermal amplification reactions is nonspecific amplification, which cannot be mitigated by thermal cycling, as in PCR. Investigators have employed a range of techniques to mitigate this pitfall, including engineered thermally stabilized polymerases and two-step nested reactions targeting different regions of the amplicon of interest (Nested Mango NASBA/NMN). These strategies, while they enable high sensitivity and specificity, necessarily add complexity to the reaction.^8,9^

One potential source of nonspecific amplification in reactions employing viral polymerases, such as T7 RNA polymerase, is unpromoted transcription. This latent behavior of T7 RNA polymerase was recently shown to be the reason transcripts produced by this enzyme can exhibit innate immune activation. This occurs because the enzyme produces a low level of antisense transcript, which hybridizes with the main sense transcript, producing dsRNA and causing innate immune activation.^10^ We reasoned that unpromoted transcription could be operative in NASBA reactions, generating a primer dimer that codes for the fluorescent aptamer, which can go on to seed the reaction. Here, we demonstrate that the inclusion of a competitor duplex that does not contain a promoter sequence can fully suppress nonspecific NASBA amplification. Using this technique, we demonstrate multicolor tagging of NASBA amplicons with fluorescent aptamers of diverse sequence and structure, including in multiplexed, one-pot reactions with real-time readout.

## Results

Apta-NABSA entails the use of two primers: the first is a conventional NASBA primer, which installs the T7 RNA polymerase promoter sequence. The second binds the amplicon and additionally contains the antisense sequence for a functional RNA (here, a fluorogenic aptamer). Thus, as the amount of RNA amplicon transcribed increases, the amount of RNA aptamer increases. The aptamer can then fold and bind its corresponding small molecule, generating a specific, real-time fluorescent signal (Figure 1). The use of primers coding for a fluorescent aptamer is particularly appealing, since this enables the specificity typically associated with a dual-labeled probe (e.g., TaqMan, molecular beacons), coupled with the low cost associated with a small molecule dye (e.g., SYBR Green).^2^ We designed primers for an enteroaggregative *E. coli* virulence gene, *aggR*, encoding a transcriptional activator of aggregative adherence fimbriae I using primers previously developed for real-time PCR by Hidaka, et al.^11^ The primer lacking the T7 RNA polymerase promoter sequence contained the coding sequence for the Broccoli aptamer, acting as a genetically encoded, green fluorescent RNA tag when complexed with the fluorogenic dye DFHBI-1T.^12^

**Figure 1.**
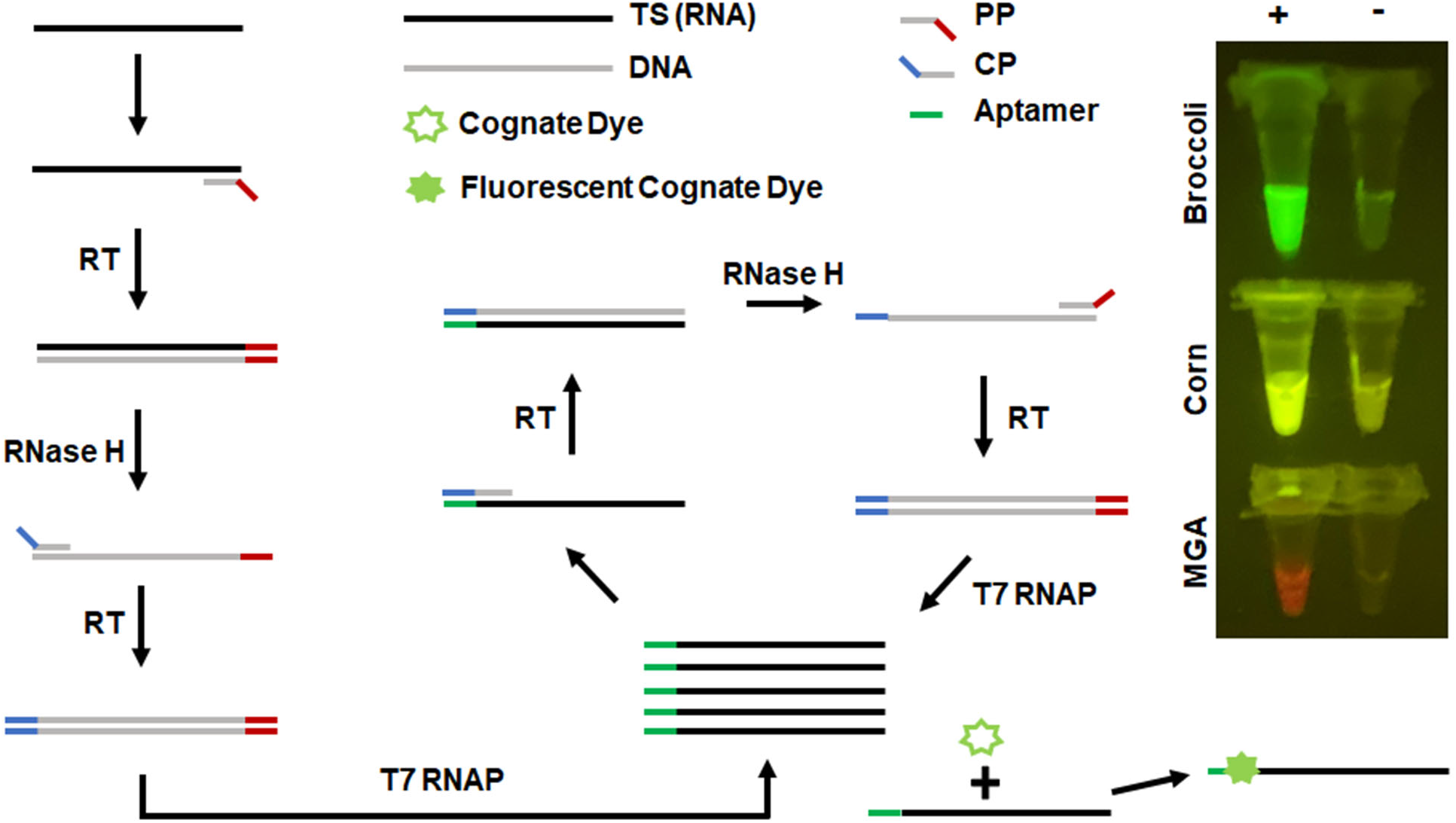
Apta-NASBA reaction. Schematic of the reaction. 1) Promoter-containing primer (**PP**) binds the amplicon of interest. Reverse transcriptase (**RT**) extends **PP**, generating a DNA-RNA hybrid comprised of the amplicon target sequence (**TS**) and an antisense DNA copy of the amplicon with an overhang containing the T7 RNA polymerase (T7 RNAP) promoter. **RNase H** degrades the original amplicon RNA of interest. Coding primer (**CP**) binds. **RT** extends **CP**, generating a DNA duplex corresponding to the amplicon, with a **T7 RNAP** promoter (red) and an aptamer coding sequence (blue). **T7 RNAP** generates an RNA fusion construct that contains the sequence antisense to **TS**, fused at the 3′ end with the aptamer corresponding to the sequence in **CP** (green). **CP** binds this RNA, which again enters the RT-RNase H-T7 RNAP cycle. At each step, RNA is generated with multiple turnover, enabling exponential amplification. Images on the right are representative Apta-NASBA reactions using aptamers that span the fluorescent spectrum.

As expected, fluorogenic aptamer-encoding primers gave exponential amplification curves (Figure 2A). However, as others have observed in NASBA reactions, untemplated amplification occurred in template-free controls (Figures 2A-2E).^8^ We speculated that unpromoted T7 RNA polymerase activity, as was recently reported,^10^ could have been the source of the observed false positives. Given that either the sense or antisense RNA strand is sufficient to seed a NASBA reaction, and that our NASBA primers contain the T7 RNA polymerase promoter in one primer and the aptamer coding sequence in the other, a small amount of primer dimer could cause a false positive.

**Figure 2.**
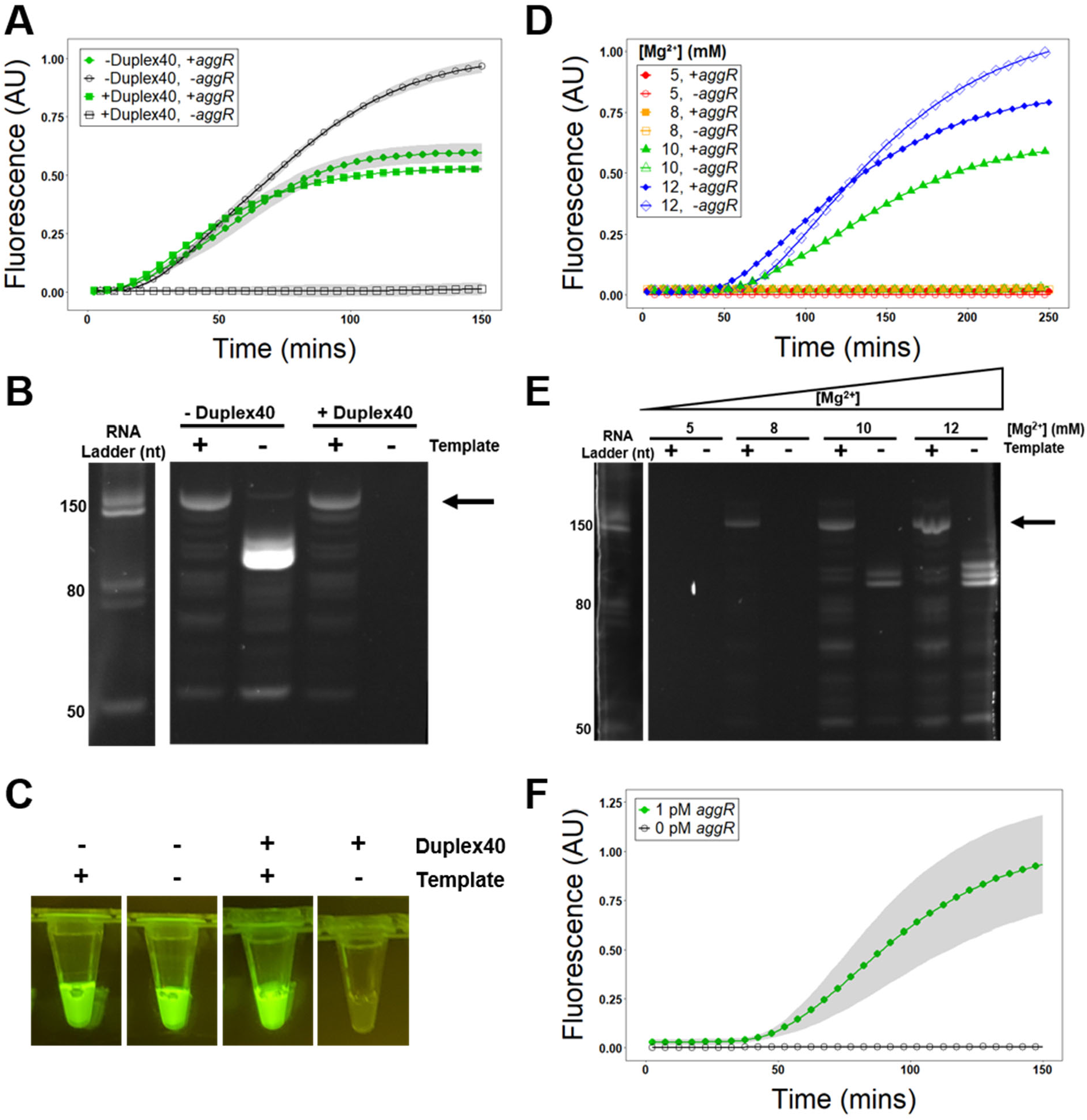
NASBA is highly prone to false positives, which can be suppressed by modulation of reaction conditions. Apta-NASBA was performed using the enteroaggregative *Escherichia coli* gene, *aggR*. Typical NASBA reactions exhibit fluorescence developing over ca. 2 h, with frequent false positives. These false positives can be suppressed by inclusion of **Duplex40** as observed by fluorescence spectroscopy (Panel **A**) and PAGE (Panel **B**). Panel **A**: samples containing template=green closed markers, samples lacking template=black open markers, samples containing **Duplex40**=circles, samples lacking **Duplex40**=squares). Panel **B**: Left: SYBR Gold stained ladder, right: DFHBI stained gel. Arrow indicates size of expected full-length amplicon-aptamer fusion product (144 nt); false positives are of lower size, consistent with a primer-derived product and not template contamination. Panel **C:** Images of Apta-NASBA reactions with and without **Duplex40.** False positives can also be suppressed by decreased Mg^2+^ concentration, which has previously been reported to suppress unpromoted transcription^10^, as observed by fluorescence spectroscopy (Panel **D**) and PAGE (Panel **E**). Panel **D**: samples containing template=closed markers, samples lacking template=open markers. Samples containing 12 mM Mg^2+^ (blue diamonds) exhibited false positives, while 10 mM Mg^2+^ (green triangles) still exhibited NASBA with ca. 40% reduction in signal intensity, and false positives were not detected by fluorescence spectroscopy, but they were detected by PAGE. Samples containing 8 mM Mg^2+^ (orange squares) did not exhibit detectable product by fluorescence spectroscopy, but product was detectable by PAGE. 5 mM Mg^2+^ did not support NASBA amplification (red circles). Panel **E**: Left: SYBR Gold stained ladder, right: DFHBI stained gel. Arrow indicates size of expected full-length amplicon-aptamer fusion product (144 nt); false positives are of lower size, indicating a primer-derived product, and not template contamination. Panel **F**: Optimized Apta-NASBA enables specific sensitive detection of *E. coli* virulence gene *aggR* at an amplicon concentration as low as 1 pM.

We reasoned that the presence of a competitor duplex of a sequence unrelated to the amplicon of interest and lacking both the T7 RNA polymerase promoter and the aptamer-encoding sequence could suppress this phenomenon. We screened a range of concentrations of a 40 bp duplex **Duplex40** (**Supplementary Figure S1**) and observed that **Duplex40**, when present at 10 μM duplex, fully suppressed untemplated amplification with a minimal diminution in final fluorescence intensity relative to reactions lacking **Duplex40** (Figures 2A-2C). A longer, 100 bp duplex **Duplex100** also sufficed to suppress false positives, with similar effectiveness at a lower concentration - 0.316 μM duplex (**Supplementary Figure S2**).

The optimum concentration of competitor duplex DNA (in base pairs) decreased with increasing length, with an optimum of 400 μM bp for **Duplex40** and 31.7 μM bp for **Duplex100**. We thus speculated that T7 RNA polymerase could be associated with nucleic acids for a duration of time in proportion to their length, and that very long nucleic acids could act as ultra-potent inhibitors of false positives. However, this was not the case; genomic (calf thymus) DNA did not suppress false positives (**Supplementary Figure S3**). We also screened heparin, another highly negatively charged polyelectrolyte and inhibitor of T7 RNA polymerase, at submaximal levels using a Spinach-tagged Apta-NASBA construct (**Supplementary Figure S4**).^13–15^ Heparin reduced the positive and false positive signals by approximately the same extent.

We also screened Mg^2+^ concentrations, previously reported to suppress unpromoted transcription.^10^ Decreasing Mg^2+^ from 12 mM to 10 mM diminished false positive signal, albeit with a decrease in signal in positive reactions (Figure 2D). A further decrease to 8 mM fully suppressed unpromoted transcription with a further diminution in final fluorescence intensity (Figure 2D) resulting in the positive reaction only being detectable by a weak band on PAGE (Figure 2E).

We performed further optimization of the reaction using **Duplex40** as the false-positive inhibitor. Optimizations included removing chloride (**Supplementary Figure S5**),^16^ altering primer concentration (**Supplementary Figure S6**), adding osmolytes (**Figures S7-S12**), altering potassium concentration (**Supplementary Figure S13**), and adding inorganic pyrophosphatase (**Supplementary Figure S14**).^17^ Optimized reaction conditions resulted in a limit of detection of 1 pM (Figure 2F), with no signal from negative reactions (i.e., those lacking template).

Having observed that the Apta-NASBA system functions with Broccoli, a green aptamer (λ_ex_=472 nm, λ_em_=507 nm), we sought to expand Apta-NASBA into different colors, using Corn, a yellow aptamer (λ_ex_=505 nm, λ_em_=545 nm).^18,19^ We designed primers for two additional pathogen-associated genes, *estP* (observed using Corn) and *estH* (observed using Broccoli) by modifying primers previously developed for real-time PCR.^22^ *estP* and *estH* produce two different variants of heat-stable enterotoxins found in enterotoxigenic *E. coli*, and is a major cause of diarrhea in travelers and infants.^20–22^ Additionally, we designed another primer for *aggR* to observe using malachite green aptamer, a red aptamer (λ_ex_=630 nm, λ_em_=650 nm).^20,23,24^ Reactions were successful using Corn to detect *estP* and malachite green aptamer to detect *aggR* (Figure 3A and **Supplementary Figure S15**). Consistent with the high degree of specificity we found **Duplex40** to afford, amplicons only exhibited signal when amplified with their cognate primer set (**Supplementary Figure S16**).

**Figure 3.**
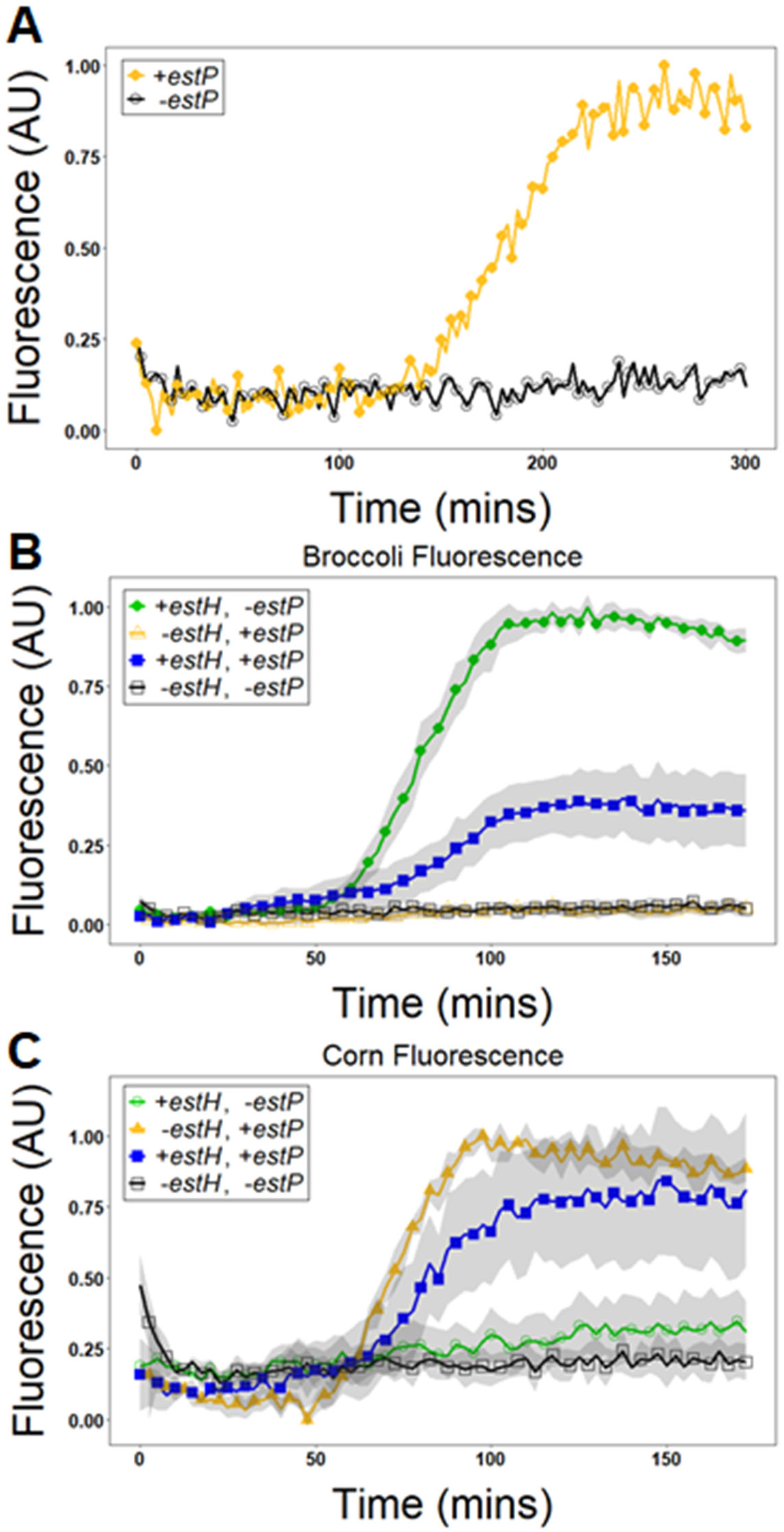
Apta-NASBA functions with multiple fluorescent aptamers. Panel **A**: Apta-NASBA of enteroinvasive *E. coli estP* gene with Corn (λ_ex_= 505 nm, λ_em_= 545 nm) detecting 218 pM template RNA performed with SpectraMax Gemini XS plate reader. The ability to use different fluorescent aptamers allows for multiplexing. Multiplexed Apta-NASBA reactions using Broccoli to detect *E.coli estH* gene and *E. coli estP* gene with Corn (Panels **B** and **C**). Panel **B**: Multiplexed Apta-NASBA reactions detected using Broccoli. Panel **C**: Multiplexed Apta-NASBA reactions detected using Corn. Samples where signal should be seen = closed symbols, samples where no signal should be seen = open signal. Green symbols represent reactions containing template being detected with Broccoli, blue symbols represent reactions containing both template detected with Broccoli and template detected with Corn, yellow symbols represent reactions containing template detected with Corn, black symbols represent reactions containing no template (Panels **B** and **C**).

Given the strict primer-to-amplicon relationship, we sought to use these orthogonal primer pairs in a multiplexed detection system. Here we show detection of *estP* and *estH* in a single reaction using Corn and Broccoli, respectively (Figure 3B, 3C). We have also demonstrated dual-detection *estP* and *aggR* using combinations of Broccoli, Corn and the malachite green aptamer for read-out (**Supplementary Figures S17** and **S18**).

The isothermal nature of Apta-NASBA eliminates the need for energy intensive thermocyclers. To further Apta-NASBA’s utility in field diagnostics and low resource areas we coupled a lyophilized Apta-NASBA reaction with a low-cost Raspberry Pi-based detection platform employing a cell phone camera module (Figure 4C). A short Python script was written to control the image acquisition of fluorescent reactions and fluorescent intensity was determined based on pixel value.^25^ The fluorescent curves obtained from Apta-NASBA on more expensive plate reader and qPCR-based fluorescent detection platforms are functionally identical to those observed on the Raspberry Pi-based detection platform (Figure 4A-4C). We show, with the addition of trehalose, Apta-NASBA reactions are viable post lyophilization, allowing for transportation to remote locations without a cold chain (Figure 4C and **Supplementary Figure S19** and **Supplementary Video S1**).

**Figure 4.**
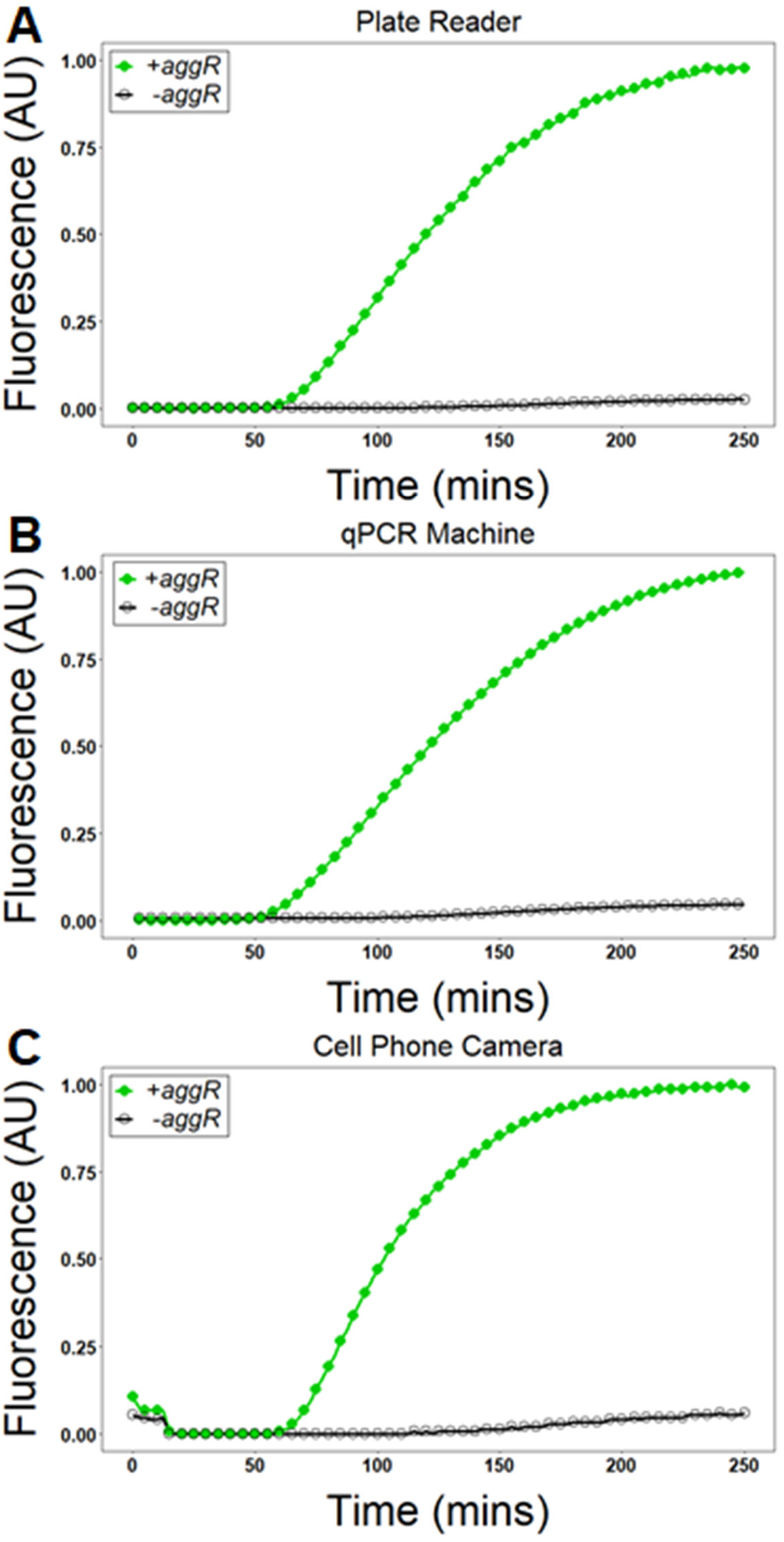
Apta-NASBA is able to be transported to remote environments using lyophilization, and Raspberry Pi-based fluorescent imaging apparatus. Panel **A**: Lyophilized Apta-NASBA reaction of *aggR* gene performed in a fluorescent plate reader (SpectraMax Gemini XS). Panel **B:** Lyophilized Apta-NASBA reaction of *aggR* gene performed in qPCR machine (ChaiBio Open qPCR). Panel **C:** Lyophilized Apta-NASBA reaction of *aggR* gene performed with the Raspberry Pi-based fluorescent imaging apparatus. All detection platforms show a characteristic template-dependent sigmoidal amplification curve.

The computer-aided design of multidomain RNAs containing fluorescent RNAs can be problematic, given that a significant number of known fluorescent RNAs fold into G-Quadruplex structures. We observed that, despite predicting an incorrect fold, the use of folding prediction software was generally successful in predicting whether a domain of a given RNA would fold independently, or induce misfolding (**Supplementary Figure S20**).

The RNA constructs developed in this paper provide proof-of-concept for design of multidomain RNAs that exhibit diverse folds, including those not accommodated by secondary structure prediction algorithms – particularly G-Quadruplexes (**Supplementary Figure S20**).^26^ These appear to be ubiquitous in fluorescent aptamers selected *in vitro*. For example, folding prediction algorithms showed Spinach, the first published G-Quadruplex-containing fluorescent RNA aptamer, to be a stem-loop structure.^15^ However, Spinach is a G-Quadruplex not captured by existing folding prediction algorithms.^27,28^ The presence of G-Quadruplexes in Apta-NASBA readouts is the reasoning behind an increase in potassium leading to a greater signal. Monovalent cations are required for proper G-Quadruplex folding and stability (**Supplementary Figure S13**). Diverse and novel folds have also been observed in several other fluorescent aptamers. The work in this paper provides important proof of concept for design and deployment of arbitrary RNAs, including those forming non-canonical secondary structures not accounted for by sequence prediction software.

Apta-NASBA represents an attractive platform for pathogen detection with the capability to be performed in remote, low resource areas. Given its modularity and compatibility with one-pot, multicolor sequence detection, it promises to be a highly extensible platform. In the current work, the readout employed is a fluorescent aptamer. Given that the functional RNA is installed using the second primer, the Apta-NASBA platform is highly versatile, and the possibility exists to install arbitrary nucleic acids, including other functional RNAs, such as aptamers for modulation of a cell-free protein expression system, or Small Transcription Activating RNAs (STARs).

## Materials and Methods

Oligonucleotides were obtained from Integrated DNA Technologies desalted and used as received. Enzymes were from New England Biolabs (Ipswitch, MA), except for T7 RNAP and mMuLV, which were overexpressed in-house as described below.

### NASBA Reaction

NASBA was performed based on extensive modifications to a procedure originally reported by Sooknanan, van Gemen, and Malek.^29^ Reactions were run with a total volume of 25 µL. A 5 X Buffer stock was prepared containing 40 mM Tris-Acetate, 84 mM KOAc, 12 mM MgOAc, 1 mM each of dATP, dTTP, dGTP and dCTP, 2 mM each of ATP, UTP, GTP and CTP. An enzyme mix was prepared containing T7 RNAP, Moloney Murine Leukemia Virus reverse transcriptase (mMuLV), RNase H (New England Biolabs Inc., M0297) and bovine serum albumin (New England Biolabs Inc., B9000S). 5X Buffer was added to a final concentration of 1 X to a solution containing 1 µM Primer 1, 1 µM Primer 2, 100 µM cognate dye, 1 mM DTT, 166 mM KOAc, 10 µM **Duplex40**, 400 mM trehalose, 1 X inorganic pyrophosphatase (Bayou Biolabs) as described previously^30^ and 4 mM MgOAc. Reactions were heated for 5 minutes at 65 °C followed by 5 minutes at 40 °C. Enzyme mix was then added to the reaction and reactions were incubated at 40 °C. For lyophilized reactions, the full reaction except amplicon and dye was flash-frozen in liquid nitrogen and lyophilized overnight using a LABCONCO FreeZone 1 liter benchtop freeze dry system, then resuspended in a solution of amplicon and dye. No 65 °C step was performed for lyophilized reactions. Reactions without addition of template were used as negative controls. Fluorescence spectroscopy has been normalized in each graph by subtracting the smallest value from each measurement followed by dividing by the largest value. Template used was transcribed *in vitro*, see below. Primer sequences can be found in Supplementary Table S1. Primers were adapted from previously published primers from Hidaka, et al.^11^ Geneious 10.1.3 was used to assist in primer design.

### T7 RNAP and mMuLV-RT Overexpression and Purification

10 mL LB containing 100 µg/µL carbenicillin were inoculated with *E. coli* DH5α containing either pT7-911Q (T7 RNAP)^31^ or pET-MRT (mMuLV-RT)^32^. Cultures were grown overnight at 37 °C, then used to inoculate an additional 1 L of LB containing 100 µg/µl carbenicillin and grown at 37 °C to an OD600 between 0.5 and 1. Cultures were then induced with 1 mM IPTG, and grown at 37 °C for 3 h. Culture was cooled on ice for 20 mins and pelleted at 3700 RPM for 15 mins. Pellets were flash frozen in liquid nitrogen and frozen at −80 °C overnight. Pellets were held in a cold room for 30 mins, then dissolved in 20 mL lysis buffer (50 mM HEPES-KOH pH 7.6, 1 M NH_4_Cl, 10 mM MgCl_2_, 7 mM BME). Pellets were incubated in lysis buffer for 30 mins followed by tip sonication. Sonication was performed at 50% power in 15 s intervals until 2 kJ total energy had been applied, then the sample was allowed to cool for 5 mins. This was repeated a total of 4 times. Pellets were then centrifuged for 45 min at 15 000 x *g* at 4 °C.

The supernatant was applied to 0.6 mL Ni-NTA agarose beads (GoldBio, H-350-50) and incubated on a rocker in a cold room for 1 h. Washing and elution steps were done in batch method. Beads were washed with 10 mL wash buffer for 10 mins then washed again with 10 mL wash buffer (50 mM HEPES pH 7.6, 1 M NH_4_Cl, 10 mM MgCl_2_, 15 mM imidazole, 7 mM BME) for 15 mins. 3 mL elution buffer (50 mM HEPES-KOH pH 7.6, 100 mM KCl, 10 mM MgCl_2_, 300 mM imidazole, 7 mM BME) was applied to beads and incubated on a rocker for 12 mins in a 4 °C cold room. Elution was dialyzed against 500 mL 2 X storage buffer (100 mM Tris-HCl pH 7.6, 200 mM KCl, 20 mM MgCl_2_, 14 mM BME) using Slide-A-Lyzer Dialysis Cassette, 2000 MWCO (Thermo Fisher Scientific, 66203) overnight, followed by dialysis against an additional 500 mL 2 X storage buffer for 3 h. Enzymes intended for lyophilization were prepared in the same storage buffer with the omission of glycerol.

Proteins were quantified using the calculated A_280_ using a NanoDrop ND-1000. Protein activity was assessed by *in vitro* transcription of Broccoli aptamer and kinetic monitoring on a fluorescence plate reader (T7 RNAP) or reverse transcription of *estH* gene and visualization on a SYBR Gold-stained urea-PAGE gel (mMuLV-RT).

### Template Transcription

1 mL transcriptions were performed to obtain template for NASBA reactions. Reactions contained 40 mM Tris-acetate, 24 mM Mg-glutamate, 100 mM K-glutamate, 2 mM spermidine, 1 mM DTT, 4 mM ATP, 4 mM GTP, 4 mM UTP, 4 mM CTP, 1 µM overexpressed T7 RNA polymerase (conversely New England Biolabs T7 RNA Polymerase M0251L can be used, but it does not give as great a yield), 1X Pyrophosphatase as used in Heili et al. and double stranded DNA templates were used according to Supplementary Table S1.^21^ Optimal template concentration was determined empirically and was typically ca. 100 nM for the amplicons used here. Reactions were incubated for 8 h at 37 °C. For transcript purification, reactions were first treated with 1 U/µg Turbo DNase (Thermo Fisher Scientific, S11494) for 15 mins at 37 °C followed ethanol precipitation. The precipitated RNA was resuspended in 1 X loading buffer (1 X TBE, 8 M Urea) and separated on a 10% Urea-PAGE gel; the full-length product band was excised via UV shadowing on a fluorescent TLC plate. The gel slice was physically disturbed by forcing it through a 21 Ga needle in 1 to 3 mL 1 X TBE, followed by 3 freeze-thaw cycles alternating between liquid nitrogen and a 55 °C water bath. The TBE containing the RNA was decanted, and an ethanol precipitation was performed. The final product was suspended in RNase-free water, and concentration was determined by A_260_ using a NanoDrop ND-1000.

### Calf Thymus DNA

Calf thymus DNA (Sigma D1501-100MG) was diluted in 10 mM Tris-Acetate pH 8, 1 mM EDTA to 2 mg/mL, and rotated until homogeneous. In order to minimize shearing, no vortexing was performed. To determine the amount of DNA used, solution concentration was determined to be 2.32 mM by A_260_ using a NanoDrop ND-1000.

### Gel Staining

10% Urea-PAGE gels (1 X TBE, 8 M urea) were incubated in two changes of folding buffer (100 mM Tris-Acetate pH 8, 10 mM MgCl_2_, 500 mM KCl) for 20 mins each time to remove urea from gels and to supply K^+^ and Mg^2+^ to support aptamer folding. Gels were then incubated in folding buffer spiked with 10 μM DFHBI for 15 mins. If the fluorescence background on a gel was high, it was destained for 5 mins in folding buffer. Gels were imaged on a gel doc (Omega Lum G, Aplegen) using the orange filter. For visualization of non-aptamer containing sequences, gels were stained in 1 X SYBR Gold for an additional 15 mins and re-imaged under the same conditions.

### 3D Printed Cell Phone Camera Module/Raspberry Pi Imaging Device

The Raspberry Pi imaging device and codes were used as described previously.^25^

## Acknowledgements

We thank members of the Engelhart laboratory for helpful discussions. This work was supported by NASA Contract 80NSSC18K1139 under the Center for Origin of Life (to A.E.E. and K.P.A.).

