## Supplementary Figures S1-S20, Supplementary Table S1 for "Highly specific, multiplexed isothermal pathogen detection with fluorescent aptamer readout"

for

**Supplementary Figure S1.** Apta-NASBA of *E. coli* *estH* gene with varying concentration of 40 bp duplex (**Duplex40**).

**Supplementary Figure S2.** Apta-NASBA of *E. coli estH* gene with varying concentration of 100 bp duplex (**Duplex100)**.

**Supplementary Figure S3.** Apta-NASBA of *E. coli* *estH* gene with varying concentration of calf thymus DNA.

**Supplementary Figure S4.** Apta-NASBA of Spinach-tagged HIV *gag* gene with addition of heparin.

**Supplementary Figure S5.** Apta-NASBA of *E. coli* *AggR* gene with chloride (■) and without chloride (●).

**Supplementary Figure S6.** Apta-NASBA of *E. coli* *estH* gene with varying primer concentrations.

**Supplementary Figure S7.** Apta-NASBA of *E. coli* *aggR* with addition of ethylene glycol.

**Supplementary Figure S8.** Apta-NASBA of *E. coli* *aggR* with addition of trehalose.

**Supplementary Figure S9.** Apta-NASBA of *E. coli* *aggR* with addition of sorbitol.

**Supplementary Figure S10.** Apta-NASBA of *E. coli* *aggR* with addition of dimethyl sulfone.

**Supplementary Figure S11.** Apta-NASBA of *E. coli* *aggR* with addition of betaine.

**Supplementary Figure S12.** Overlaid comparison of optimal concentration of each osmolyte for Apta-NASBA of *E. coli* *aggR* gene.

**Supplementary Figure S13.** Apta-NASBA of *E. coli* *estH* gene (A) Optimizing KCl concentration and (B) Optimizing KOAc concentration.

**Supplementary Figure S14.** Apta-NASBA detecting *E. coli estH* gene demonstrating the impact of inorganic pyrophosphatase on sample fluorescence.

**Supplementary Figure S15.** Apta-NASBA of *aggR* gene using the malachite green aptamer (MGA) for detection.

**Supplementary Figure S16.** Apta-NASBA does not exhibit crosstalk between primer pairs.

**Supplementary Figure S17.** Dual Apta-NASBA of *aggR* and *estP* genes using Broccoli and the malachite green aptamer (MGA), respectively, for detection.

**Supplementary Figure S18.** Dual Apta-NASBA of *aggR* and *estP* genes using the malachite green aptamer (MGA) and Corn, respectively, for detection.

**Supplementary Figure S19.** Lyophilized Apta-NASBA of *aggR* gene without trehalose (circles, ●) or with trehalose (squares, ■).

**Supplementary Video S1.** Time-lapse video of Apta-NASBA reaction collected using 3D printed Raspberry Pi-based device.

**Supplementary Figure S20.** Predicted folds of Apta-NASBA products compared to functionality in fluorescence readout assays.

**Supplementary Table S1.** Sequences of oligonucleotides used in this manuscript.

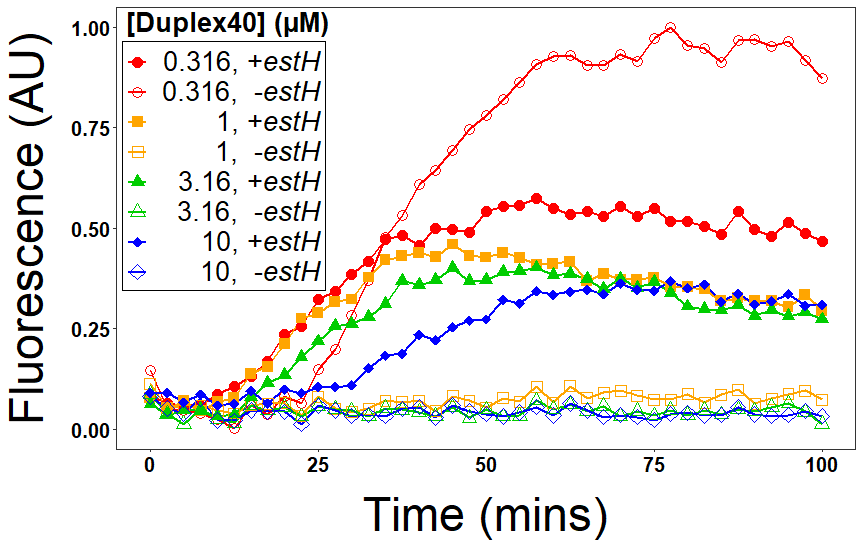
**Supplementary Figure S1.** Apta-NASBA of *E. coli* *estH* gene with varying concentration of 40 bp duplex (**Duplex40**). Positive reactions (i.e., containing amplicon) are shown using filled markers and negative reactions (i.e., lacking amplicon) are shown using open markers. As the concentration of **Duplex40** increases, the false positive signal is diminished. At 10 µM **Duplex40,** the false positive signal is fully suppressed with little decrease in the true positive signal. Greater concentrations of **Duplex40** have a negative effect on the sensitivity of the reaction.

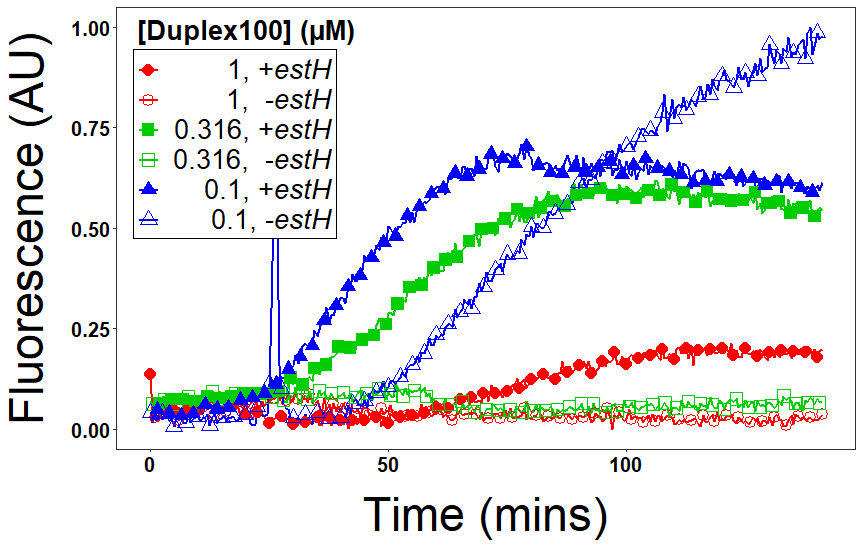

**Supplementary Figure S2.** Apta-NASBA of *E. coli estH* gene with varying concentration of 100 bp duplex (**Duplex100)**. Similar to the addition of the 40 bp duplex, as the concentration of 100 bp duplex (**Duplex100**) increases, the non-templated, false positive signal decreases. A smaller concentration of **Duplex100** compared to **Duplex40** is needed to equally suppress the false positive signal.

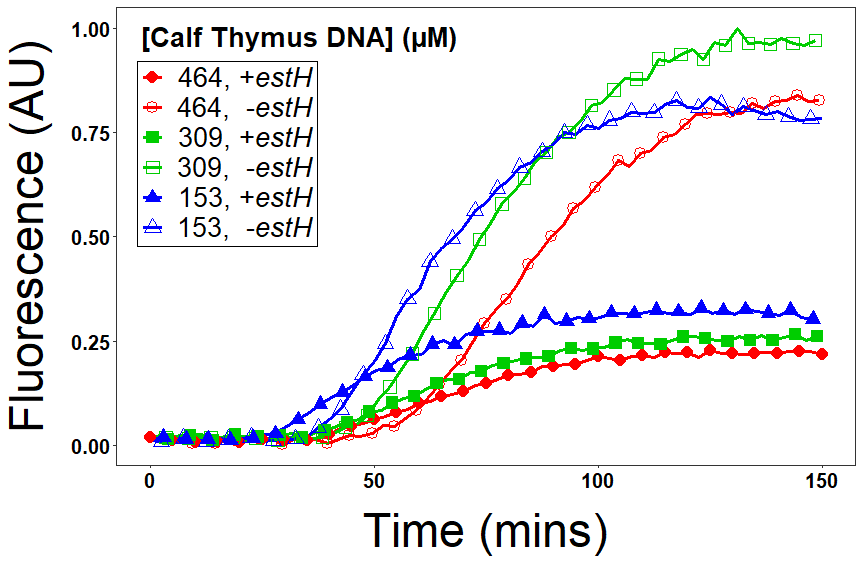
**Supplementary Figure S3.** Apta-NASBA of *E. coli* *estH* gene with varying concentration of calf thymus DNA. Positive reactions (i.e., containing amplicon) are shown using filled markers and negative reactions (i.e., lacking amplicon) are shown using open markers.

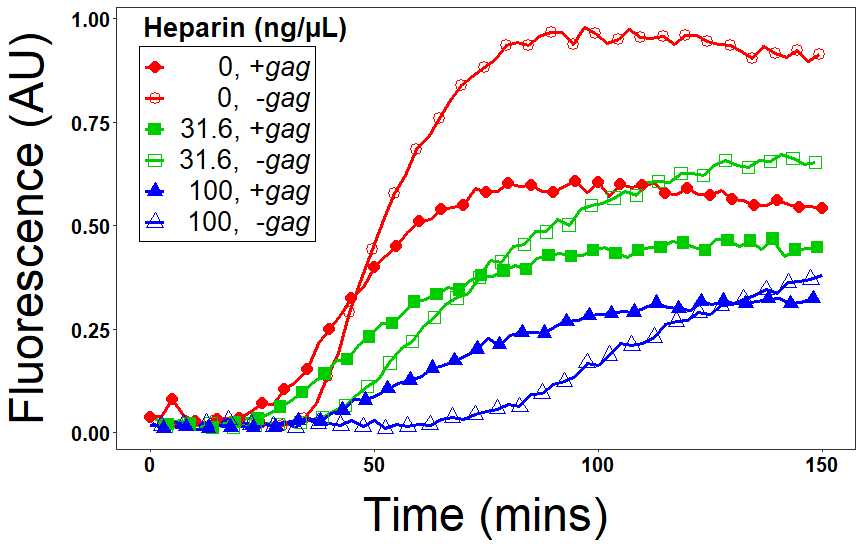

**Supplementary Figure S4.** Apta-NASBA of Spinach-tagged HIV *gag* gene with addition of heparin. Positive reactions (i.e., containing amplicon) are shown using filled markers and negative reactions (i.e., lacking amplicon) are shown using open markers.

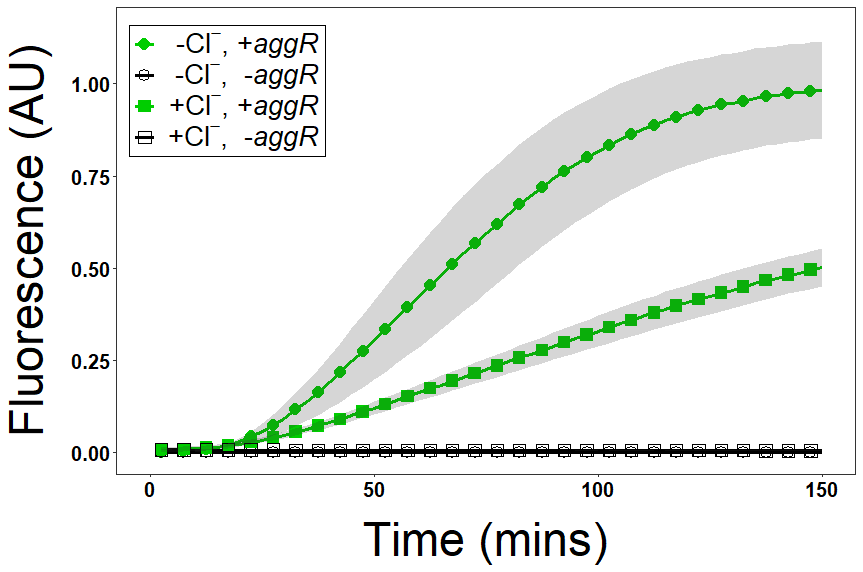

**Supplementary Figure S5.** Apta-NASBA of *E. coli* *AggR* gene with chloride (■) and without chloride (●). Positive reactions (i.e., containing amplicon) are shown using filled markers and negative reactions (i.e., lacking amplicon) are shown using open markers.

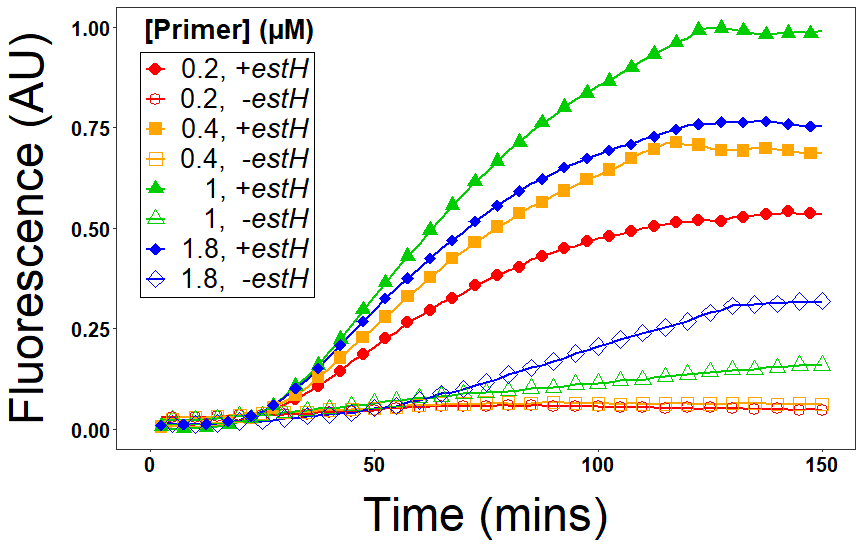

**Supplementary Figure S6.** Apta-NASBA of *E. coli* *estH* gene with varying primer concentrations. Positive reactions (i.e., containing amplicon) are shown using filled markers and negative reactions (i.e., lacking amplicon) are shown using open markers.

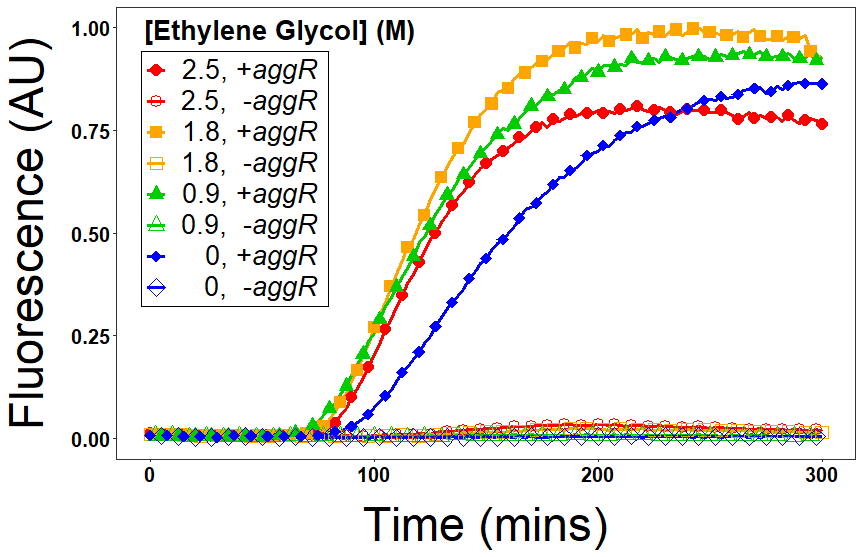

**Supplementary Figure S7.** Apta-NASBA of *E. coli* *aggR* with addition of ethylene glycol. Positive reactions (i.e., containing amplicon) are shown using filled markers and negative reactions (i.e., lacking amplicon) are shown using open markers.

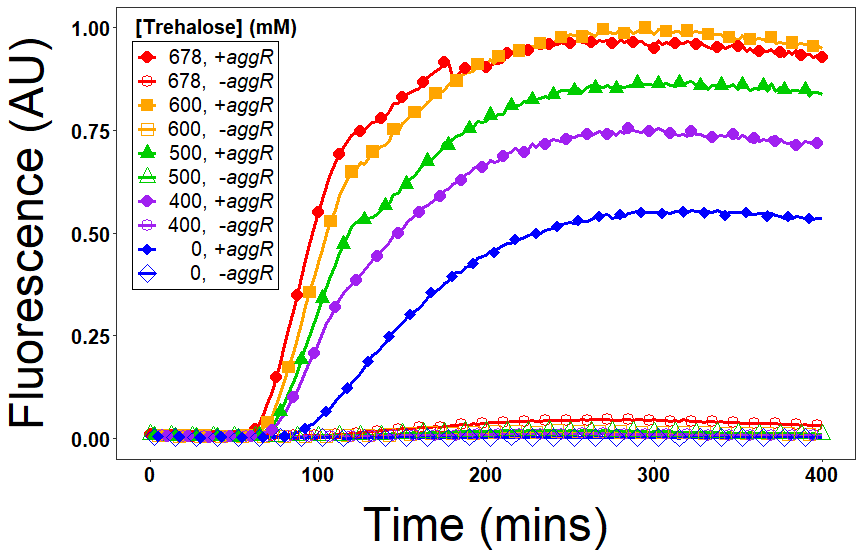

**Supplementary Figure S8.** Apta-NASBA of *E. coli* *aggR* with addition of trehalose. Positive reactions (i.e., containing amplicon) are shown using filled markers and negative reactions (i.e., lacking amplicon) are shown using open markers.

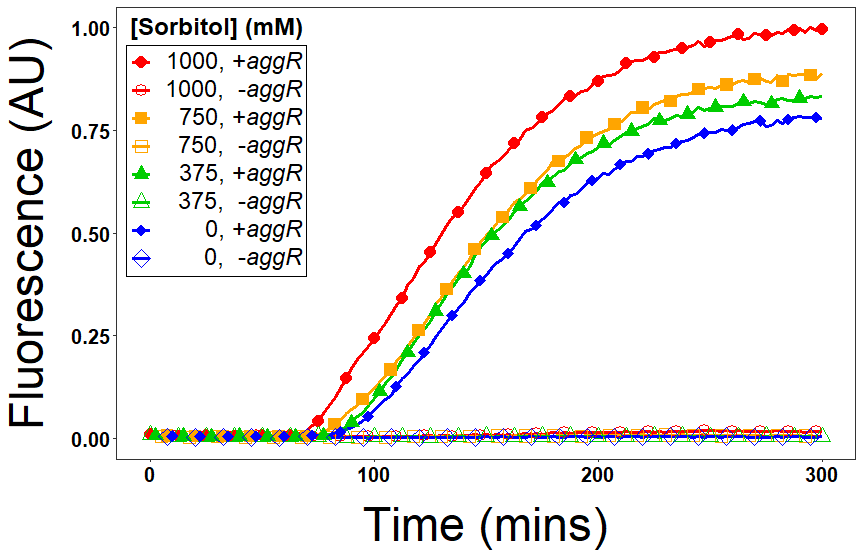

**Supplementary Figure S9.** Apta-NASBA of *E. coli* *aggR* with addition of sorbitol. Positive reactions (i.e., containing amplicon) are shown using filled markers and negative reactions (i.e., lacking amplicon) are shown using open markers.

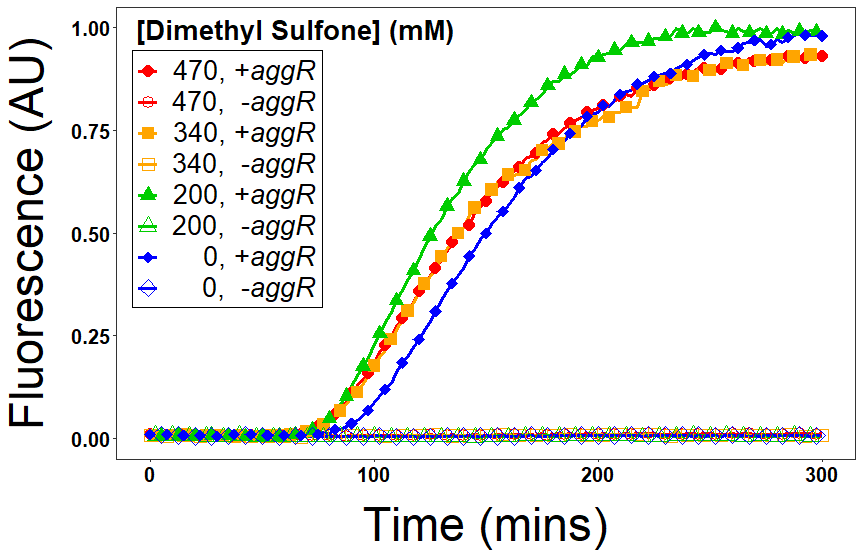
**Supplementary Figure S10.** Apta-NASBA of *E. coli* *aggR* with addition of dimethyl sulfone. Positive reactions (i.e., containing amplicon) are shown using filled markers and negative reactions (i.e., lacking amplicon) are shown using open markers.

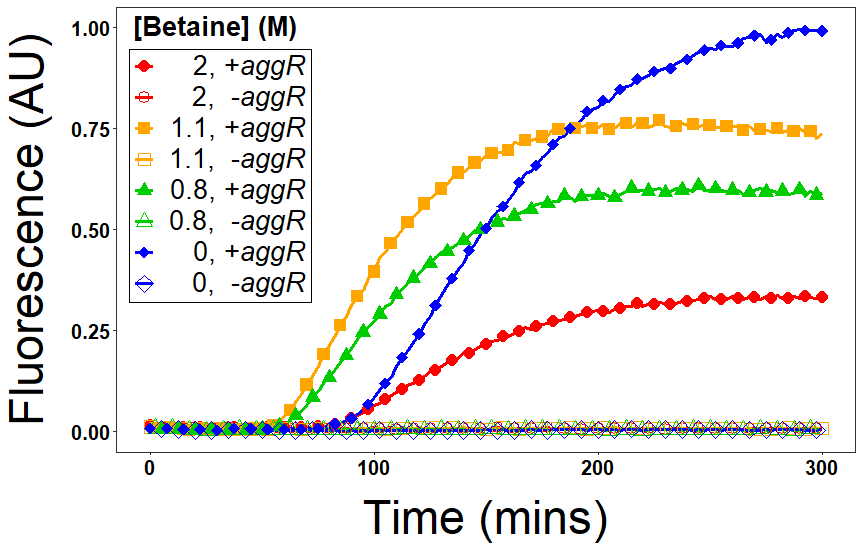
**Supplementary Figure S11.** Apta-NASBA of *E. coli* *aggR* with addition of betaine. Positive reactions (i.e., containing amplicon) are shown using filled markers and negative reactions (i.e., lacking amplicon) are shown using open markers.

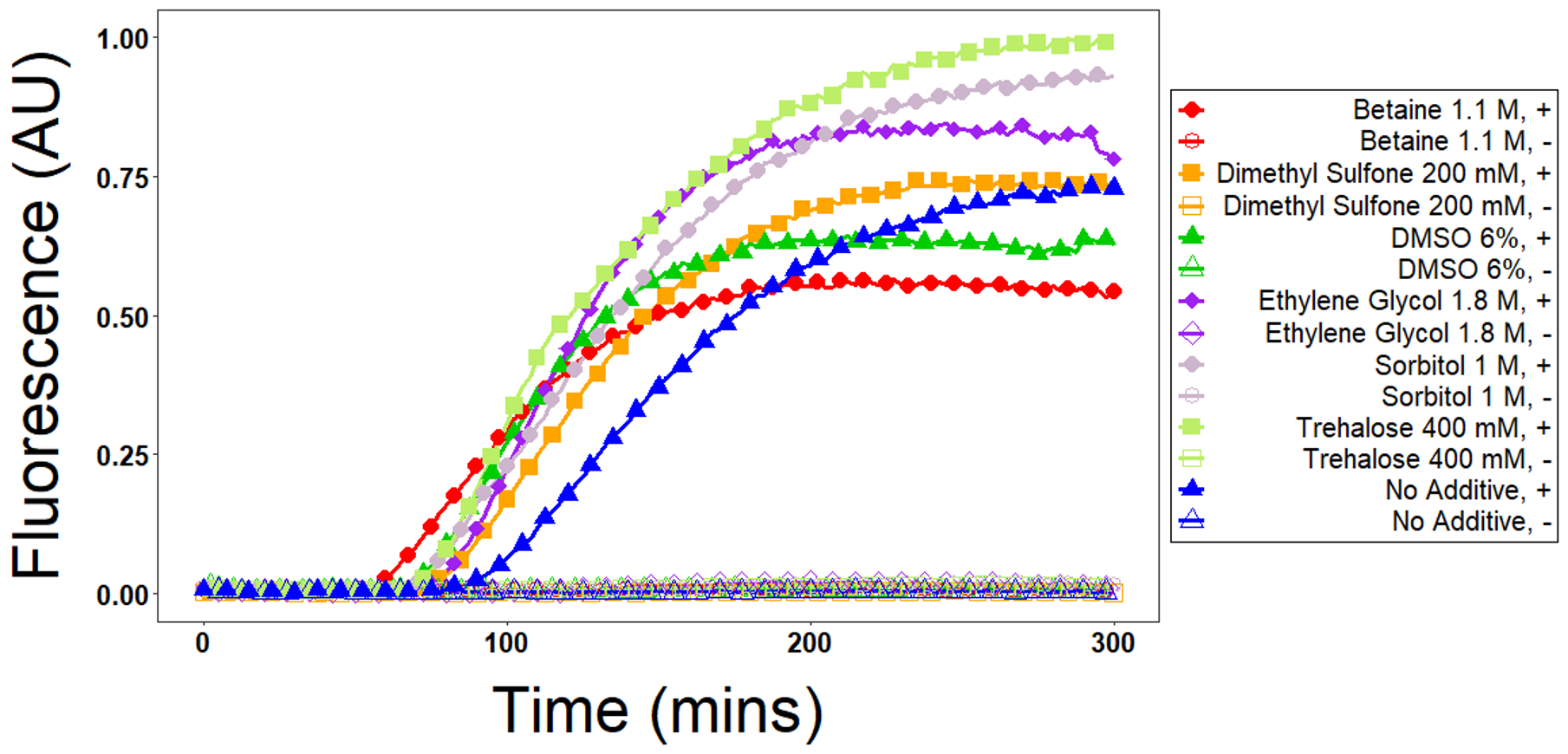

**Supplementary Figure S12.** Overlaid comparison of optimal concentration of each osmolyte for Apta-NASBA of *E. coli* *aggR* gene. Positive reactions (i.e., containing amplicon) are shown using filled markers and negative reactions (i.e., lacking amplicon) are shown using open markers.

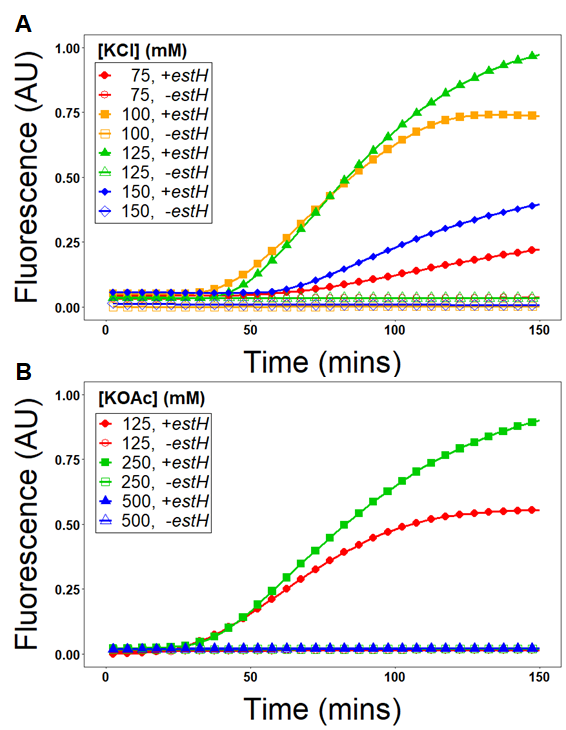

**Supplementary Figure S13.** Apta-NASBA of *E. coli* *estH* gene (A) Optimizing KCl concentration and (B) Optimizing KOAc concentration. When present as the acetate salt, the optimum concentration of potassium was higher (250 mM) than when potassium was present as the chloride salt (125 mM), likely due to elimination of inhibition of T7 RNAP by chloride. Positive reactions (i.e., containing amplicon) are shown using filled markers and negative reactions (i.e., lacking amplicon) are shown using open markers.

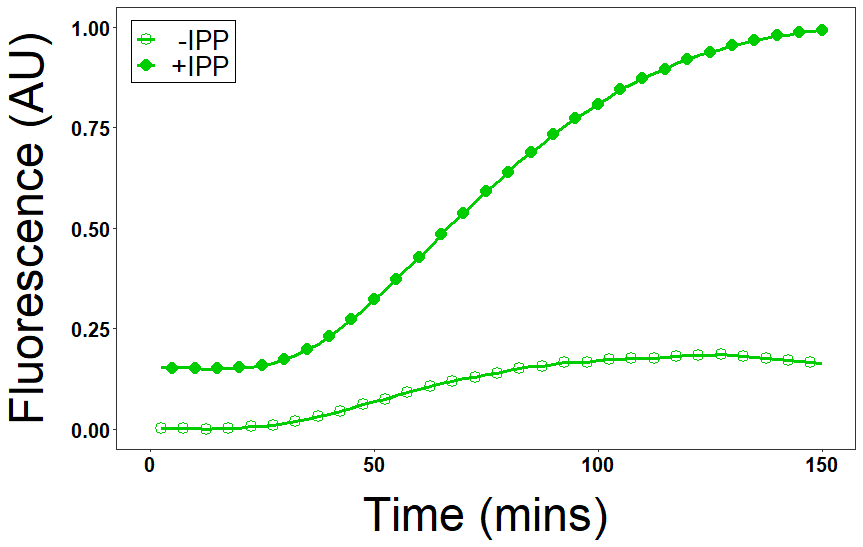

**Supplementary Figure S14.** Apta-NASBA detecting *E. coli estH* gene demonstrating the impact of inorganic pyrophosphatase on sample fluorescence.

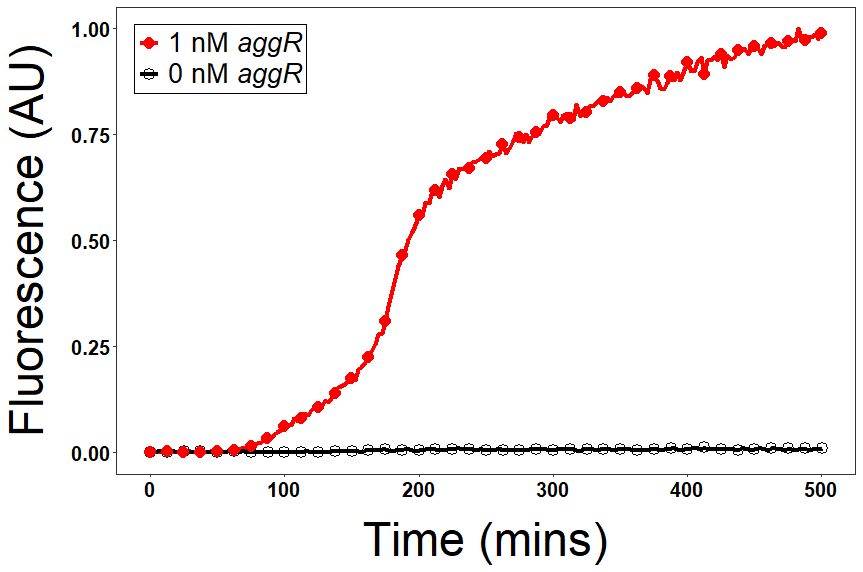

**Supplementary Figure S15.** Apta-NASBA of *aggR* gene using the malachite green aptamer (MGA) for detection.

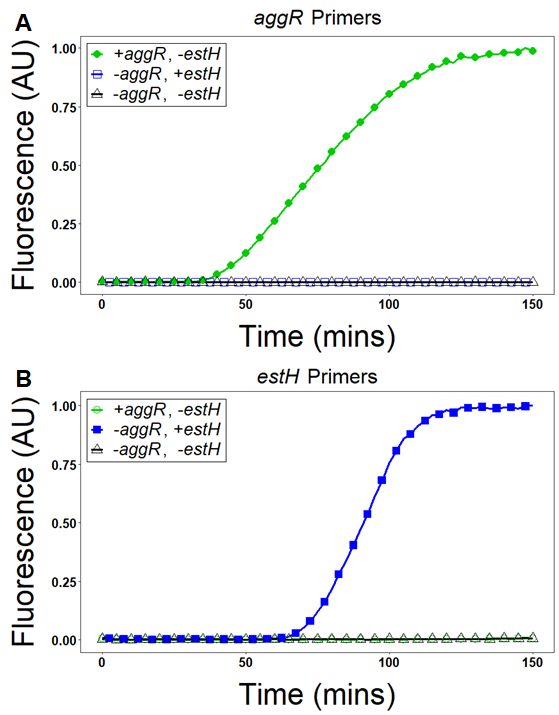
**Supplementary Figure S16.** Apta-NASBA does not exhibit crosstalk between primer pairs. Apta-NASBA of reactions containing *aggR* or *estH* produces fluorescent signal only when the amplicon and its cognate primers are present.

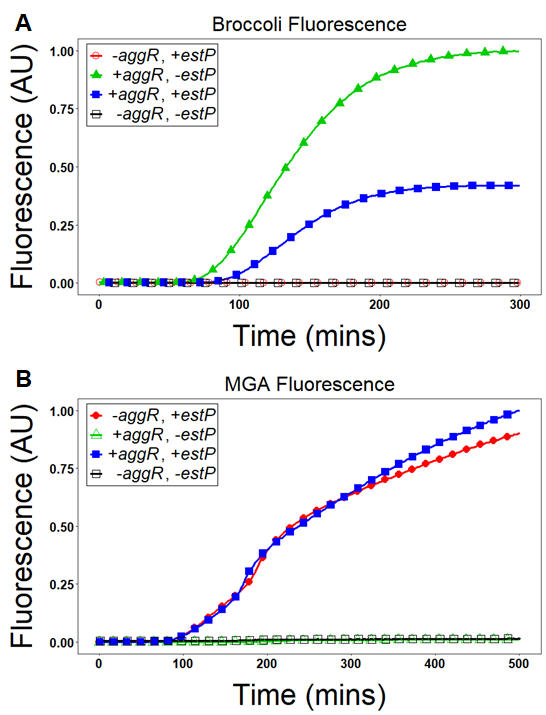

**Supplementary Figure S17.** Dual Apta-NASBA of *aggR* and *estP* genes using Broccoli and the malachite green aptamer (MGA), respectively, for detection.

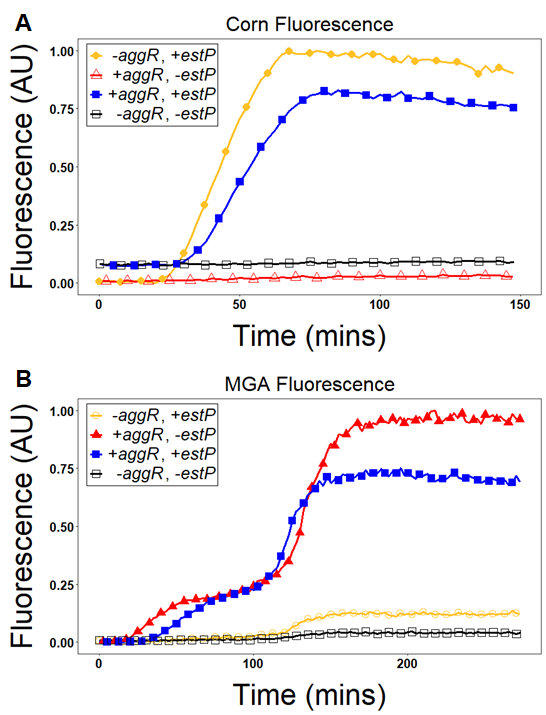

**Supplementary Figure S18.** Dual Apta-NASBA of *aggR* and *estP* genes using the malachite green aptamer (MGA) and Corn, respectively, for detection.

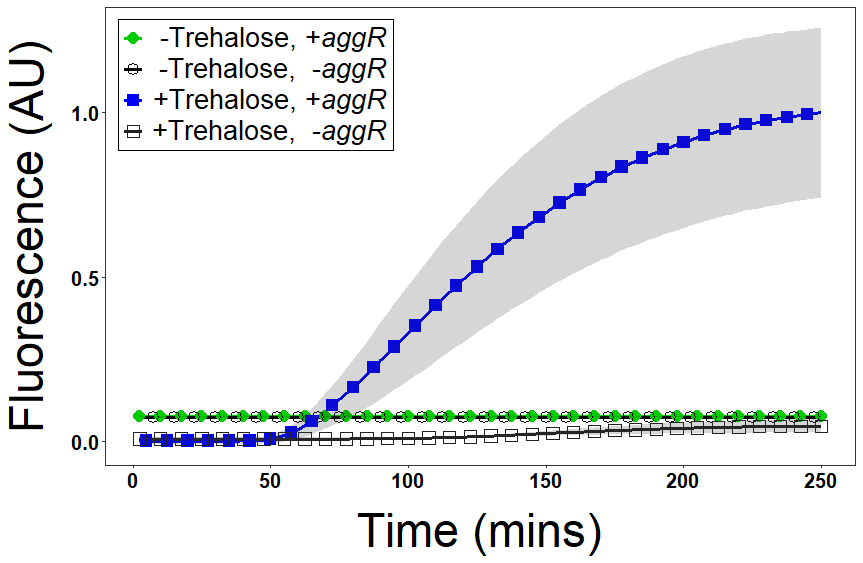
**Supplementary Figure S19.** Lyophilized Apta-NASBA of *aggR* gene without trehalose (circles, ●) or with trehalose (squares, ■). Positive reactions (i.e., containing amplicon) are shown using filled markers and negative reactions (i.e., lacking amplicon) are shown using open markers.

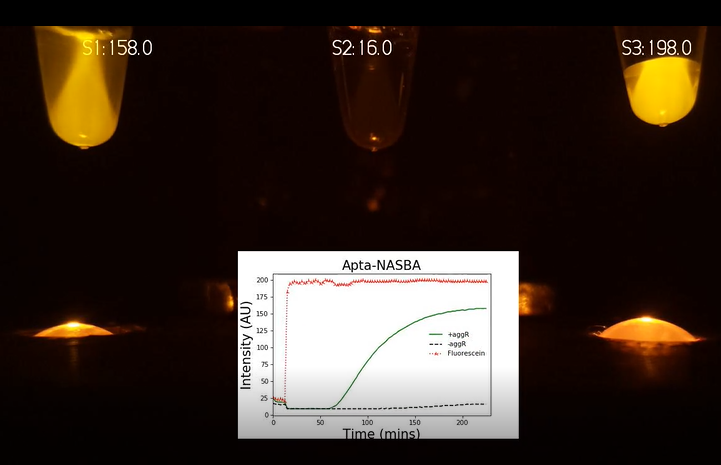
**Supplementary Video S1.** Time-lapse video of Apta-NASBA reaction collected using 3D printed Raspberry Pi-based device. Tubes, left to right: positive sample, negative sample, fluorescein standard. Still from video shown.

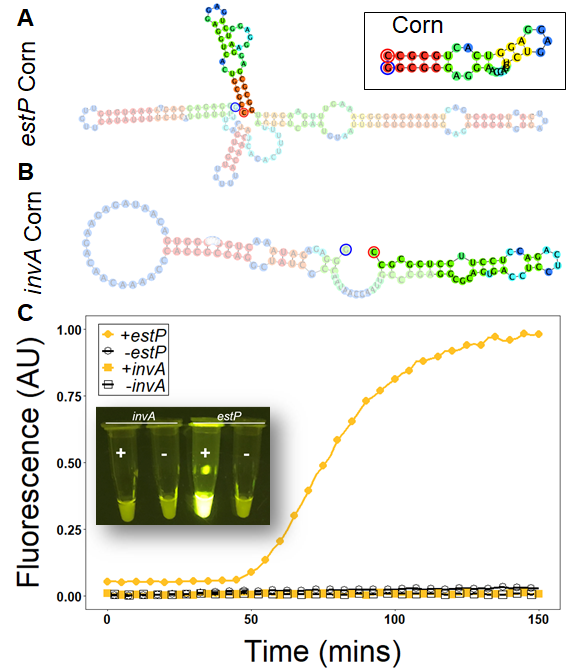

**Supplementary Figure S20.** Predicted folds of Apta-NASBA products compared to functionality in fluorescence readout assays. Panel **A:** Folding prediction of Apta-NASBA product for *estP* gene detected by Corn using Geneious 10.1.3. Product corresponding to Broccoli is highlighted and product corresponding to target amplicon is lightened. Inset shows Corn is predicted to fold similarly in the amplicon fusion context as to how it folds independently. Panel **B**: Panel **A:** Folding prediction of Apta-NASBA product for *invA* gene detected by Corn using Geneious 10.1.3. Product corresponding to Broccoli is highlighted and product corresponding to target amplicon is lightened. Inset shows Corn is predicted to interact with amplicon sequence and not fold correctly. Panel **C:** Fluorescence time course showing *estP-*Corn, but not *invA-*Corn, functions in Apta-NASBA. Inset contains fluorescence images of reactions showing same.

| **Name** | **Sequence (5′-3′)** |
| --- | --- |
| **Templates for amplicon transcription** | |
| *aggR* gene with **T7 promoter** | **AAT TCT AAT ACG ACT CAC TAT AGG GAG A**CA GCG ATA CAT TAA GAC GCC TAA AGG ATG CCC TGA TGA TAA TAT ACG GAA TAT CAA AAG TAG ATG CTT GCA GTT GTC CGA ATT GGT CAA AAG GAA TAA TTG TAG CTG ATG CTG ACG |
| *aggR* forward PCR primer | AAT TCT AAT ACG ACT CAC TAT AGG |
| *aggR* reverse PCR primer | CGT CAG CAT CAG CTA CAA TTA |
| *estP* gene with **T7 promoter** | **AAT TCT AAT ACG ACT CAC TAT AGG GAG A**CT TTC CCC TCT TTT AGT CAG TCA ACT GAA TCA CTT GAC TCT TCA AAA GAG AAA ATT ACA TTA GAG ACT AAA AAG TGT GAT GTT GTA AAA AAC AAC AGT GAA AAA AAA TCA GAA AAT ATG AAC AAC ACA TTT TAC TGC |
| *estP* forward PCR primer | AAT TCT AAT ACG ACT CAC TAT AGG |
| *estP* reverse PCR primer | GCA GTA AAA TGT GTT GTT CAT AT |
| *estH* gene with **T7 promoter** | **TAA TAC GAC TCA CTA TAG G**CA CCT TTC GCT CAG GAT GCT AAA CCA GTA GAG TCT TCA AAA GAA AAA ATC ACA CTA GAA TCA AAA AAA TGT AAC ATT GCA AAA AAA AGT AAT AAA AGT GAT CCT GAA AGC ATG AAT AGT AGC AAT TAC TG |
| *estH* forward PCR primer | TAA TAC GAC TCA CTA TAG GCA CC |
| *estH* reverse PCR primer | CAG TAA TTG CTA CTA TTC ATG CTT TCA G |
| *gag* gene with **T7 promoter** | **TAA TAC GAC TCA CTA TAG G**TG CTA AAC ACA GTG GGG GGA CAT CAA GCA GCC ATG CAA ATG TTA AAA GAG ACC ATC AAT GAG GAA GCT GCA GAA TGG GAT AGA GTG CAT CCA GTG CAT GCA GGG CCT ATT GCA CCA GGC CAG ATG AGA GAA CCA AGG GGA AGT GAC ATA GCA |
| *gag* forward PCR primer | TAA TAC GAC TCA CTA TAG GTG CTA AAC A |
| *gag* reverse PCR primer | TGC TAT GTC ACT TCC CCT TGG |
| *invA* gene with **T7 promoter** | **AAT TCT AAT ACG ACT CAC TAT AGG GAG A**TC GGG CAA TTC GTT ATT GGC GAT AGC CTG GCG GTG GGT TTT GTT GTC TTC TCT ATT GTC ACC GTG GTC CAG TTT ATC |
| *invA* forward PCR primer | AAT TCT AAT ACG ACT CAC TAT AGG |
| *invA* reverse PCR primer | GAT AAA CTG GAC CAC GGT GA |
| **NASBA** **primers** | |
| *aggR* **T7** primer | **AAT TCT AAT ACG ACT CAC TAT AGG GAG A**CG TCA GCA TCA GCT ACA ATT ATT CC |
| *aggR* **Broccoli** primer | **GAG CCC ACA CTC TAC TCG ACA GAT ACG AAT ATC TGG ACC CGA CCG TCT C**CA GCG ATA CAT TAA GAC GCC TAA AG |
| *aggR* **MGA** Primer | **GGA TCC ATT CGT TAC CTG GCT CTC GCC AGT CGG GAT** CAG CGA TAC ATT AAG ACG CCT AAA G |
| *estP* **T7** primer | **AAT TCT AAT ACG ACT CAC TAT AGG GAG A**GC AGT AAA ATG TGT TGT TCA TAT TTT CTG |
| *estP* **MGA** primer | **GGA TCC ATT CGT TAC CTG GCT CTC GCC AGT CGG GAT** CTT TCC CCT CTT TTA GTC AGT CAA CT |
| *estP* **Corn** primer | **GGC GCA GTG ACC TCC TCA GAC CTC CTT CCT CGC GCC** AAT CTT CAA ACT TTC CCC TCT TTT AGT CAG TCA ACT |
| *estH* **T7** primer | **AAT TCT AAT ACG ACT CAC TAT AGG GAG A**CA GTA ATT GCT ACT ATT CAT GCT TTC AG |
| *estH* **Broccoli** primer | **GAG CCC ACA CTC TAC TCG ACA GAT ACG AAT ATC TGG ACC CGA CCG TCT C**CC TTT CGC TCA GGA TGC TAA AC |
| *gag* **T7** primer | **AAT TCT AAT ACG ACT CAC TAT AGG G**TG CTA TGT CAC TTC CCC TTG GTT CTC TCA |
| *gag* **Spinach** primer | GCT ACA TTC GAA TCA ACA A**GA CGC GAC CAG TTA CGG AGC TCA CAC TCT ACT CAA CAG TGC CGA AGC ACT GGA CCC GTC CTT CAC CAT TTC GGT CGC GTC** CCT CTA CGC TTA AGA CTG CAC CAG TTT CGT CCT CAC GGA CTC ATC AGA CCG GAA AGC ACA TCC GGT AGT GGG GGG ACA TCA AGC AGC CAT GCA AA |
| *invA* **T7** primer | **AAT TCT AAT ACG ACT CAC TAT AGG GAG A**GA TAA ACT GGA CCA CGG TGA CA |
| *invA* **Corn** primer | **GGC GCG AGG AAG GAG GTC TGA GGA GGT CAC TGC GCC** TCG GGC AAT TCG TTA TTG G |
| **Inhibitor duplexes** | |
| Duplex100 | GGT GTC AGT AAG CCA TTC GAG ATC CTC ATA GTC GTC TCA CCA GAT CCT CAT AGT CGT CTC ACC AGA TCC TCA TAG TCG TCT CAC CAG ATC CCT TGG GTT G (top)  CAA CCC AAG GGA TCT GGT GAG ACG ACT ATG AGG ATC TGG TGA GAC GAC TAT GAG GAT CTG GTG AGA CGA CTA TGA GGA TCT CGA ATG GCT TAC TGA CAC C (bottom) |
| Duplex40 | GGT GTC AGT AAG CCA TTC GAG ATC CTC ATA GTC GTC TCA C (top)  GTG AGA CGA CTA TGA GGA TCT CGA ATG GCT TAC TGA CAC C (bottom) |

**Supplementary Table S1.** Sequences of oligonucleotides used in this manuscript.
